# Hippocampal signature of associative memory measured by chronic ambulatory intracranial EEG

**DOI:** 10.1101/291740

**Authors:** Simon Henin, Anita Shankar, Nicolas Hasulak, Daniel Friedman, Patricia Dugan, Lucia Melloni, Adeen Flinker, Cansu Sarac, May Fang, Werner Doyle, Thomas Tcheng, Orrin Devinsky, Lila Davachi, Anli Liu

**Affiliations:** New York University Comprehensive Epilepsy Center, 223 34^th^ Street, New York, NY 10016, USA; Department of Neurology, New York University School of Medicine, 240 East 38^th^ Street, 20^th^ Floor, New York, NY 10016, USA; NeuroPace, Inc., 455 N. Bernardo Avenue, Mountain View, CA 94043, USA.; Department of Neuroscience, Max Planck Institute for Empirical Aesthetics, Gruneburgweg 14, 60322 Frankfurt am Main, Germany.; Department of Psychology, New York University, 6 Washington Place, New York, NY 10003, USA.; Department of Neurosurgery, New York University School of Medicine, 530 1^st^ Avenue, Suite 7W, New York NY 10016, USA; Columbia University, Department of Psychology, 1190 Amsterdam Ave #406, New York, NY, 10027, USA.

## Abstract

Some patients with medically refractory focal epilepsy are chronically implanted with a brain-responsive neurostimulation device (the RNS^®^ System), permitting neurophysiological measurements at millisecond resolution. This clinical device can be adapted to measure hippocampal dynamics time-locked to cognitive tasks. We illustrate the technique with a proof of concept in three patients previously implanted with the RNS System as they engage in an associative memory task, measured months apart. Hippocampal activity measured in successful encoding in RNS System patients mirrors that in surgical patients during intracranial electroencephalography (iEEG), suggesting that chronic iEEG allows sensitive measurements of hippocampal physiology over prolonged timescales.

---

Conventional intracranial electroencephalography (iEEG) monitoring may be performed during evaluation for resective epilepsy surgery and is the gold standard for fine-grained dissection of the spatiotemporal dynamics of cognitive processing. Compared to noninvasive methods, such as scalp EEG or functional magnetic resonance imaging (fMRI), iEEG recordings offer an excellent signal-to-noise ratio and direct measurements of neuronal responses at millisecond resolution. However, limitations to task-based research utilizing iEEG in the hospital setting include: (1) Suboptimal patient participation due to clinical factors including seizures, pain, medications, fatigue, disrupted sleep-wake cycles, and post-surgical recovery and (2) Lack of control over the hospital environment which may cause distraction during testing^1^. These clinical considerations affect subject participation as well as the quality of the iEEG data. Sources of environmental noise, such as 60 Hz line noise generated from nearby equipment (e.g. hospital bed, compression stockings, and monitoring equipment) and frequent interictal epileptiform discharges must be handled during signal analysis, resulting in significant data loss. Furthermore, the duration of iEEG is limited to 3-16 days due to infection risk and degradation of signal quality. Therefore, cognitive research utilizing this method is limited to testing within single sessions, or at best repeated sessions over days^2^. For these reasons, questions regarding the longitudinal nature of memory processing have been difficult to study.

Some patients with refractory focal epilepsy are implanted with a brain-responsive neurostimulation device (RNS^®^ System, NeuroPace, Inc.), which presents an opportunity to study memory dynamics and other cognitive functions under more controlled experimental conditions and over prolonged time scales. The RNS System is an FDA-approved therapy to treat medically refractory focal seizures in adults^3, 4^. At detection of abnormal electrical activity, the RNS System delivers brief pulses of electrical stimulation, programmed by the physician, intended to terminate seizure activity. The system includes two four-contact leads placed on or in seizure foci, which record and store changes in local-field potentials (LFP) with millisecond precision. While the RNS System is a clinical therapy, the availability of chronic electrocorticography permits the investigation of long-term brain dynamics in the ambulatory setting. For example, evoked responses to phonetic features of speech perception and production have been demonstrated to be stable over 1.5 years^5^, and circadian and ultradian patterns of long term seizure dynamics have been revealed^6–8^. Because the RNS^®^ Neurostimulator is battery powered and cranially implanted, it can record iEEG with much less environmental artifact compared to conventional iEEG (further technical characteristics in the Methods Section).

To illustrate the potential of the RNS System as a research tool, we demonstrate a method to interface with the clinical device for task-based cognitive testing. In this IRB approved study, three patients previously implanted with the RNS System consented to engage in a cross-modal associative memory task, performed during testing sessions spanning several months. Real-time iEEG recording and integration with a testing platform was enabled by IRB-approved Research Accessories (RAs), permitting task-based study design and analysis (**Figure 1**). These adults with drug-resistant focal onset epilepsy were implanted with the RNS System in accordance with FDA-approved indication for use and for reasons completely independent of this research study. Patients who were selected for participation had a cranially implanted brain-responsive programmable neurostimulator connected to hippocampal-placed depth leads and were capable of consent. As a comparison, we performed the same memory task and signal analysis in five (5) surgical patients implanted acutely for localization of the seizure focus with hippocampal depth leads. We describe the clinical characteristics of the eight (8) patients (3 RNS System, 5 surgical) in **Table 1** and in the Supplementary Material section.

**TABLE 1.** Subject characteristics and test performance

| Subj ID | Age | Sex | Hand | Lead/Electrode Coverage | FSIQ | Session Number | Date | # of trials | % correct | Average Block Size | # Trials Rejected |
| --- | --- | --- | --- | --- | --- | --- | --- | --- | --- | --- | --- |
| R1 | 21 | M | R | L Hipp depth;<br>L basal temp strip | 108 | 1 of 3 | May 2017 | 78 | 65.4 | 8.7 | 6 |
|  |  |  |  |  |  | 2 of 3 | October 2017 | 114 | 50.9 | 11.4 | 7 |
|  |  |  |  |  |  | 3 of 3 | January 2018 | 118 | 48.3 | 14.8 | 14 |
| R2 | 45 | M | R | L hipp depth;<br>R hipp depth | 84 | 1 of 1 | April 2017 | 36 | 19.4 | 6.0 | 0 |
| R3 | 32 | F | R | L hipp depth;<br>R hipp depth | 96 | 1 of 2 | November 2017 | 118 | 53.4 | 6.9 | 1 |
|  |  |  |  |  |  | 2 of 2 | January 2018 | 93 | 47.3 | 7.8 | 2 |
| S1 | 45 | F | R | L and R hipp depths (2+2);<br>L and R frontal, temporal, parietal, occipital strips (8+8) | 99 | 1 of 1 | January 2017 | 106 | 79.2 | 2.3 | 20 |
| S2 | 18 | M | R | L and R hipp depths (2+2);<br>L and R frontal, temporal, parietal, and occipital strips (8+8) | 91 | 1 of 1 | July 2017 | 70 | 42.9 | 8.8 | 16 |
| S3 | 37 | M | R | L and R hipp and insular depths (3+3); L and R frontal, temporal, parietal, and occipital strips (5+5) | 82 | 1 of 1 | October 2017 | 94 | 40.4 | 4.5 | 64 |
| S4 | 20 | F | R | R hipp and insular depths (5); R hemisphere (2 x 64 contact grids; 6 frontal and temporal strips); | 74 | 1 of 1 | December 2017 | 119 | 47.1 | 4.1 | 12 |
| S5 | 25 | M | R | R hipp depths; R hemisphere | 134 | 1 of 1 | February 2018 | 92 | 46.7 | 9.2 | 12 |

**Figure 1.**
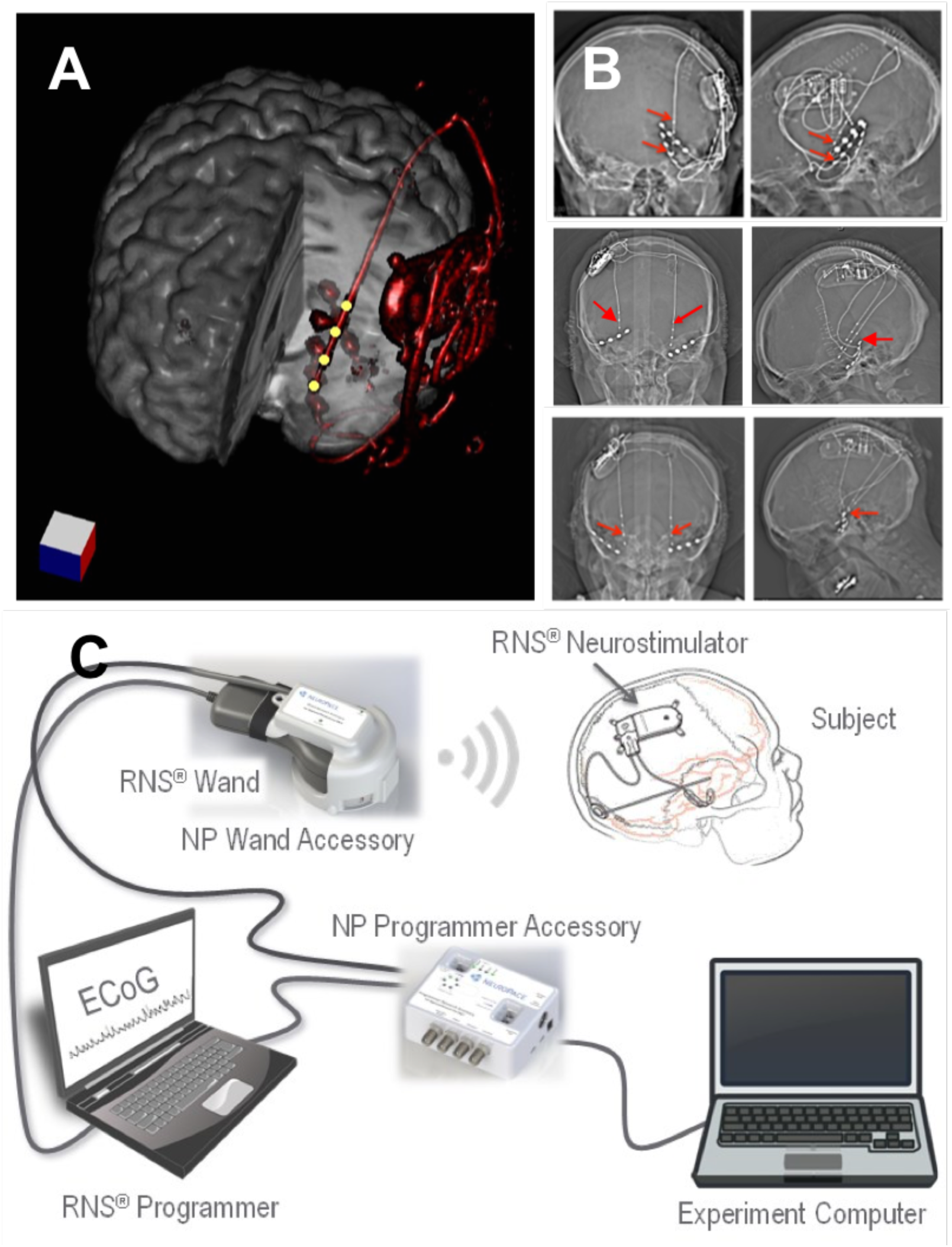
RNS System patients and experimental system. **(A)** 3-dimensional reconstruction of neurostimulator and lead placement (red). Yellow dots are recording hippocampal depth lead electrodes. **(B)** Top Row. Head CT of Patient R1 with neurostimulator and left hippocampal depth and mesial temporal subdural lead electrodes (red arrows), coronal (left) and sagittal (right) views.Additional leads present but not connected to neurostimulator, include: left basal temporal neocortical and lateral temporal neocortex. Middle and Bottom Rows. Head CT of Patient R2 and R3 with neurostimulator and bilateral hippocampal depth lead electrodes, with coronal and sagittal views. Additional leads present but not connected to neurostimulator include: bilateral basal temporal. **(C)** RNS System and Research Accessories (RAs), including Wand Accessory (WA) and Programmer Accessory (PA).

The RAs do not modify the RNS System, rather they use existing features of the clinical device while providing a computer-based method for integrating real-time markers into the ongoing recording stream (See Methods, Supplementary Figure 2), allowing for event-related neural response acquisition and analysis. The task computer was connected to the Programmer Accessory (PA) via a universal serial bus (USB) communication link, and precisely timed trigger commands were sent to the PA at stimulus onset. Because the clinical system records iEEG segments limited to a maximum of 4 minutes in duration, the task software commanded the PA to pause, store, and restart recording after intervals shorter than 4 minutes, thereby permitting continuous cognitive testing and iEEG recording. To permit sensitive detection of gamma band activity, the neurostimulator’s low-pass filter was adjusted from its default setting of 90 Hz to 120 Hz during testing and recording. Responsive stimulation was suspended during task participation to avoid interference with endogenous neural rhythms during participation. An epilepsy physician (AL, PD, DF) monitored the iEEG signal to ensure that no subclinical seizures occurred during testing, and to manage the patient in the event of a clinical seizure. Given that the RNS System iEEG is recorded in the bipolar montage between adjacent electrodes, we also analyzed the conventional EEG data using the bipolar montage (see Methods).

**Figure 2.**
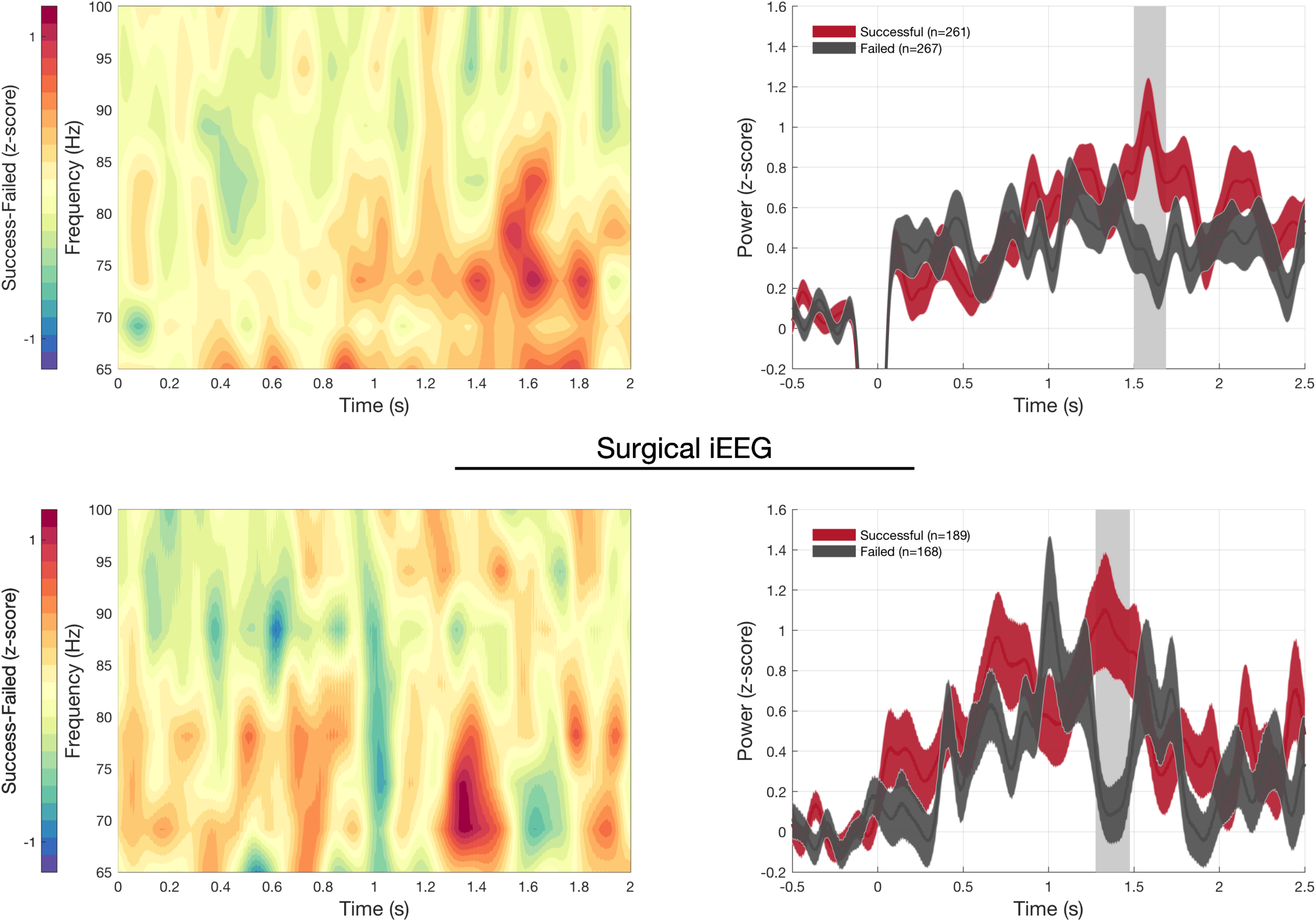
Hippocampal signature of successful versus failed encoding in RNS System and Surgical patients. Successful encoding shows increased and sustained high-gamma power in the hippocampus, compared to failed encoding in both RNS System iEEG and surgical iEEG recordings. LEFT. Differences (Successful-Failed Encoding) show increases in high-gamma power over similar frequency regions and timescales in both RNS System (top left) and surgical patients (bottom left). RIGHT: High-gamma power in hippocampal lead electrodes exhibits a similar time course, with increased high-gamma power occurring 1.25-1.6s from stimulus onset in both RNS System (top right) and surgical patients (bottom right; mean +/− SEM; shaded grey bar indicates significant differences, p<0.01 permutation test, corrected for multiple comparisons. The drop in in power from -0.1-0.05s in RNS System data is due to data blanking due to trigger artifact, see Methods).

As the hippocampus is critical to forming novel associations between previously unrelated items of information^9–11^, we utilized a cross-modal association task to examine hippocampal processing during successful associative memory encoding. During encoding, patients were shown a series of color pictures of faces paired with professions, given a brief distraction task, then were presented the same faces in a randomized order and were asked to freely recall the paired profession (see Methods). This face-profession association task has produced robust fMRI activation in bilateral hippocampi^12^. Other associative memory paradigms have produced robust activation of the anterior hippocampus on fMRI ^13–17^ (for review, see^18, 19^) with precisely localized activity to the anterior CA2 and CA3 subfields during learning of novel relationships^20^. Prior iEEG studies in surgical patients have found that gamma activity increases in the hippocampi and widespread cortical regions during encoding and can distinguish items that are later successfully remembered vs. forgotten^21–23^, termed the subsequent memory effect (SME). Increases in gamma power may represent coordinated neural firing of local cell assemblies^24^ or a more general marker of activity or stochastic volatility^25^, both of which can be meaningfully related to memory processes.

Our main finding was that increased hippocampal gamma power occurs 1.4-1.6 seconds after stimulus onset in successful versus failed encoding trials (**Figure 2**, right, p<0.01, cluster corrected permutation test, 261 successful, 267 failed trials) across our 3 subjects implanted with the RNS System. Furthermore, the hippocampal physiology supportive of successful encoding in our RNS System patients mirrored the physiological changes observed in our five (5) surgical patients in the task **(Figure 2**). When the face-profession association task was administered to five (5) surgical patients, an increase in gamma power in hippocampus occurred 1.3-1.5 seconds after stimulus presentation in successful versus failed encoding trials (**Figure 2**, p<0.01 cluster corrected permutation test, with 189 successful and 168 failed trials). Notably, there was substantial overlap in the frequency range of these gamma power increases (e.g. 65-80 Hz, **Figure 2, Left**). In addition, classification accuracy at the single trial level (e.g. prediction of successful vs. failed encoding trials based on gamma power) was above chance over the same respective time periods in both surgical and RNS System patients (**Supplemental Figure S3**).

Performance on the face-profession association task for RNS System patients varied (range 19.4 - 65.4% correct, mean 50.3 +/− 15.2 SD correct). (**Table 1**), as did performance by the surgical patients (range 40.4-79.2% correct, mean 52.2 +/− 15.9 SD correct). During outpatient behavioral testing with our RNS System patients, we found our three RNS System patients were generally able to engage in memory testing for longer periods of time (as demonstrated by the larger block sizes, **Table 1**) compared to our surgical patients. Although responsive neurostimulation was temporarily disabled, there were no seizures or other complications during testing.

Importantly, during signal analysis, we found that we discarded less iEEG data due to environmental noise or epileptiform activity in our RNS System patients compared to our surgical patients. Overall, approximately 25% (124/481) of trials were rejected from the surgical iEEG data, whereas only 5% (30/558) of the trials were rejected from the RNS System iEEG data (**Table 1**).

In summary, patients with chronically implanted brain-responsive neurostimulation devices (RNS System) present an opportunity to perform memory and other cognitive testing under more controlled conditions and across longer timescales compared to those typically used with conventional iEEG. We have developed a methodology that integrates the RNS System clinical device with a testing platform. The technique incorporates event markers into the live streaming iEEG signal from the patient’s RNS System, which allows for later task-based iEEG analysis. As a proof of principle, we have demonstrated that the hippocampal signature of successful encoding during a paired association task--a robust gamma peak at approximately 1.5 seconds after stimulus presentation--is similar in RNS System patients compared to surgical patients with hippocampal depth electrodes. While this study was performed in only a small number of patients, we have found that RNS System patients are able to engage in the memory task for longer sessions compared to our surgical patients (**Table 1**). Testing sessions in our RNS System patients were performed over the span of several months, which is not possible for the surgical patients. Furthermore, during signal analysis, fewer segments were excluded due to environmental noise or interictal epileptiform activity in the RNS System data compared to the surgical data (**Table 1**).

The major limitation of the RNS System as a research tool is restricted electrode coverage based on the patient’s clinical indication. While lead electrodes are placed on or in epileptogenic tissue, the decision to avoid surgery suggests that the tissue is functional. Another consideration is that the RNS System only permits recording in a bipolar montage, which may exclude physiological signals occurring over broader regions of hippocampus. In addition, the RNS System currently only allows for continuous recordings of up to 4 minutes, which limits the duration of any given testing block. However, this limit is overcome by introducing short interruptions (~15s) between blocks or trials to stop/store/restart the RNS System recording for investigational purposes. Finally, filter characteristics (i.e. high-pass filter at 4 Hz and low-pass filter at 120 Hz) attenuate functional signatures known to fall outside the band-pass filter range of the RNS System.

Nevertheless, patients with refractory focal epilepsy with depth lead electrodes in the hippocampus or other regions of interest offer a remarkable opportunity to study brain dynamics within more meaningful contexts. Because the neurostimulator is chronically implanted and battery powered, investigations of neurophysiology during real-world activities are now possible. Finally, the RNS System used in conjunction with Research Accessories as a research platform offers the potential of investigations conducted across more behaviorally relevant timescales (hours to days to months to years), compared to those typically used in research (seconds to minutes).

## METHODS

### Human Subjects

This study was performed in epilepsy patients with previously implanted RNS Systems for treatment of refractory focal epilepsy. Research with RNS System patients was performed in accordance with protocol approved by the New York University School of Medicine Institutional Review Board (NYUMC), and with written patient consent. Subjects were eligible if they (1) were adults (18-70 years), (2) had the RNS System with at least one hippocampal depth lead, (3) had an FSIQ>70, (4) were able to provide informed consent, and (5) were Native English speakers.

We also collected data from a group of patients undergoing surgical evaluation with intracranial EEG monitoring at NYUMC. The protocol was approved by the NYUMC Institutional Review Board and the Clinical Trials Registration number was NCT02263274 (www.clinicaltrials.gov). Subjects were eligible according to pre-established criteria, including: (1) age over 18 years old; (2) undergoing invasive monitoring for seizure localization for epilepsy surgery; and (3) ability to provide informed consent or have a legal guardian who could consent. Exclusion criteria included (1) significant cognitive impairment (IQ<70), (2) facial or forehead skin breakdown that would interfere with surface electrode placement, (3) contraindication to MRI. All patients provided informed consent. Subjects were enrolled between January 2017 and January 2018. **Table 1** lists subject characteristics.

### RNS System and Research Platform

The RNS System is an FDA-approved medical device that provides brain-responsive (closed-loop) neurostimulation for patients with medically refractory epilepsy^3, 4^. RNS System patients were chronically implanted with the RNS System for clinical reasons completely unrelated to research. Strategies for intracranial electrodes differed across patients with respect to lead type (subdural strip and/or depth) and implantation site. The lead type selection and implantation decisions were dependent on the clinical indication. The RNS System included two four-electrode leads, with 10mm spacing (from electrode center to center), placed on or in the seizure foci. For the patients each channel recorded LFPs in a bipolar montage between adjacent electrodes. The electrocorticogram is filtered between 4-120 Hz (-3.5 dB at 4.5 and 100 Hz), and digitized at 250 Hz with 10-bit A/D resolution. When used for clinical reasons, short electrocorticographic (ECoG) recordings are stored by the neurostimulator as a result of potential electrographic seizure activity, predetermined scheduling, or a magnet swipe. In a research setting, the RNS System’s live recording mode, using a wireless Wand, is used to capture ECoG records.

To facilitate research utilizing the RNS System, several non-product tools (“Research Accessories”, RAs) were designed to enable external control of several basic RNS System functions and allow for synchronization with an experiment computer without modifying the FDA approved product. The RAs include: (1) an accessory for the Programmer laptop (Programmer Accessory, PA) and (2) an accessory for the Wand (Wand Accessory, WA). The WA consists of a custom circuit board, an MSP-430 microcontroller (Texas Instruments), an electromagnet and a telemetry coil contained in a 3D printed case that encloses the Wand. The WA is controlled by commands received from the PA. The PA consists of an Arduino Due Microcontroller (“Arduino”) running custom software and a custom printed circuit board. The PA communicates with the experiment computer via a USB control interface. The Arduino is also connected to the Programmer laptop via USB and emulates a hardware mouse and keyboard to provide control inputs to the Programmer. The Wand, while placed over the neurostimulator, allows live (real time) ECoG data to be viewed on the Programmer laptop in a continuous manner. ECoG data storage is normally limited to 4 minutes by the clinical system before it must be stored with several manual button clicks on the Programmer laptop. The first function of the RAs automates storage of the ECoG data by executing a series of mouse movements and mouse clicks on the Programmer laptop graphical user interface (GUI). This is performed by stopping a currently running real time ECoG (clicking the “Stop” button), storing it to the Programmer’s hard disk (clicking the “Store” button), and then restarting the ECoG streaming (clicking the “Start” button). This configuration allows a single USB command from the experiment computer to control ECoG storage in a repeatable manner with precise timing that would not be possible if these tasks were performed manually without the RAs. This command introduces a minimal delay (~15 seconds) between recording sessions. A second function of the RAs instructs the WA to insert a pattern of markers into the telemetry signal of the streaming ECoG data received from the neurostimulator by the Wand. This is equivalent to briefly moving the Wand away from the neurostimulator, which commonly occurs during live ECoG recording, resulting in a telemetry dropout artifact. This distinctive pattern provides a means of reliably marking (“trigger marking”) the ECoG data within 2 ECoG samples (±4 ms) of the command. This marker is later used to provide synchronization of the time-series data between the experiment computer and the stored ECoG records. By inserting these telemetry marker artifacts (“triggers”) into the real time ECoG data by using the RAs near the start and end of a 4-minute ECoG recording, we are able to account for clock differences between computers and provide reliable time synchronization between the experimental events on the experiment computer and the ECoG data.

### Post-surgical Intracranial EEG Recordings

Intracranial EEG (iEEG) was recorded from implanted subdural platinum-iridium electrodes embedded in silastic sheets (2.3 mm diameter contacts, 10 mm center-center spacing, Ad-Tech Medical Instrument, Racine, WI) or depth electrodes (1.1 mm diameter, 5-10 mm center-center spacing). The decision to implant, placement of recording electrodes, and the duration of invasive monitoring were determined solely on clinical grounds and without reference to this study. Electrodes were arranged as grid arrays (8 × 8 contacts, 10 or 5 mm center-to-center spacing), linear strips (1 × 4 to 12 contacts), or depth electrodes (1 × 8 or 12 contacts), or some combination thereof. Subdural electrodes covered extensive portions of lateral and medial frontal, parietal, occipital, and temporal cortex of the left and/or right hemisphere.

Intracranial EEG is continuously streamed to a central monitoring station connected to a 40TB storage system at NYU Langone Health Tisch Hospital. A closed-circuit TV system complements the EEG recordings. Recordings at NYU are conducted in a 16-bed telemetry ward. Within 24 hours after surgical implantation of electrodes, patients underwent a post-operative brain MRI to confirm subdural electrode placement. Electrodes were localized and mapped onto the pre- and post-implant MRI using geometric models of the electrode strips/grids and the cortical surface (Yang et al., 2012).

### Clinical iEEG Recording Equipment

Recordings from grid, strip and depth electrode arrays were made using a NicoletOne C64 clinical amplifier (Natus Neurologics, Middleton, WI), bandpass filtered from 0.16-250 Hz and digitized at 512 Hz. Intracranial EEG signals were referenced to a two-contact subdural strip facing towards the skull near the craniotomy site. A similar2-contact strip screwed to the skull was used for the instrument ground.

### Electrode Localization and Electrode Selection

Electrode localization was performed using automated processes and expert review. Pre-surgical and post-surgical T1-weighted MRIs were acquired for each patient, and the location of the electrode relative to the cortical surface was determined from coregistered MRIs following the procedure described in Yang et al.^26^. Co-registered, skull-stripped T1 images were nonlinearly registered to an MNI-152 template and electrode locations were then extracted in Montreal Neurological Institute (MNI) space (projected to the surface) using the co-registered image. A three-dimensional reconstruction of each patient’s brain was computed using FreeSurfer (http://surfer.nmr.mgh.harvard.edu). Subdural electrodes were localized to neocortical regions in Freesurfer. MRI scans were co-registered using advanced normalization tools^26^ and visualized using Mango (Multi-image Analysis GUI, Research Imaging Institute, UTHSCSA). Hippocampal depth electrode location was confirmed by expert review (AL, DF).

Localization of the RNS System lead electrodes was performed by using linear co-registration of the pre-operative, skull-stripped MRI with the post-operative high-resolution CT (using BioimageSuite v3.0). Anatomical location of each contact was determined by visual inspection of the coregistered image (DF, AL). One electrode pair was selected per participant based on the anatomical localization. For surgical patients, several electrodes may be placed in the hippocampus, however, as similar visual association tasks have shown robust activation within anterior regions of the hippocampus^15–18^, we selected the electrode pair with the best signal quality located in the anterior hippocampus. The electrode selection for each patient is provided in **Table 2**.

**Table 2.**
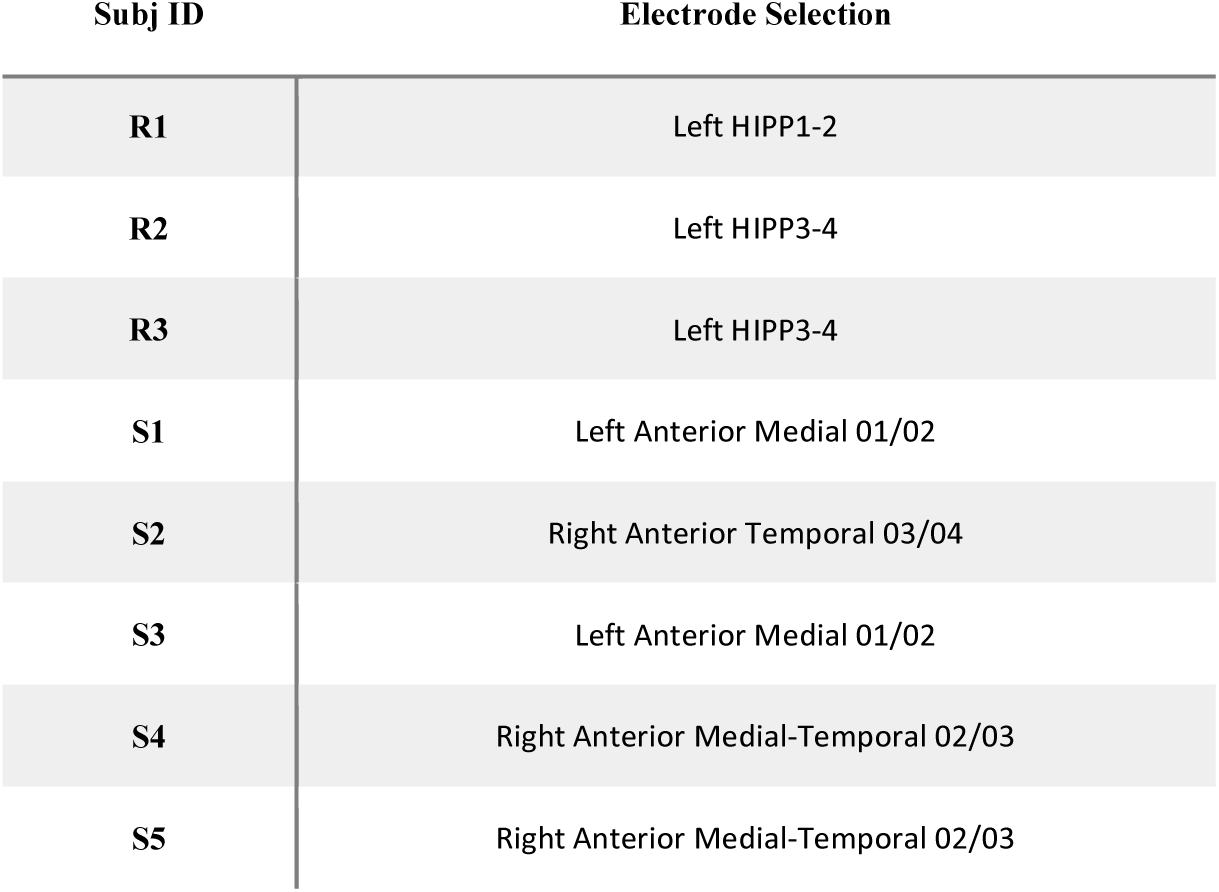
Electrode selection

**Figure. 3.**
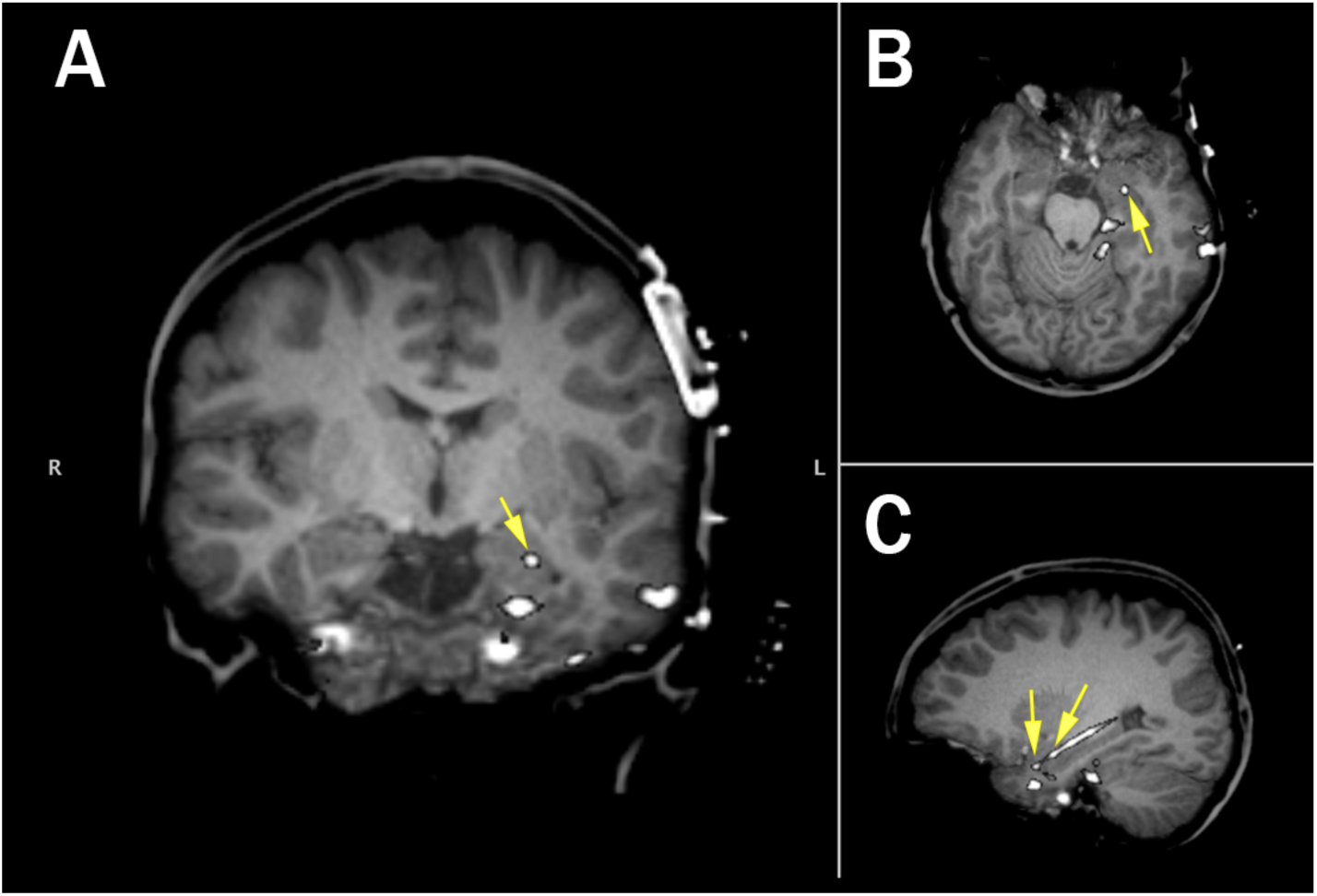
Electrode localization for RNS System patient (R1). Location of selected electrodes (Left Hipp 1-2) indicated by yellow arrows (A: Coronal, B: Axial, C: Sagittal; Coronal and Axial Views show only LH2).

**Figure 4.**
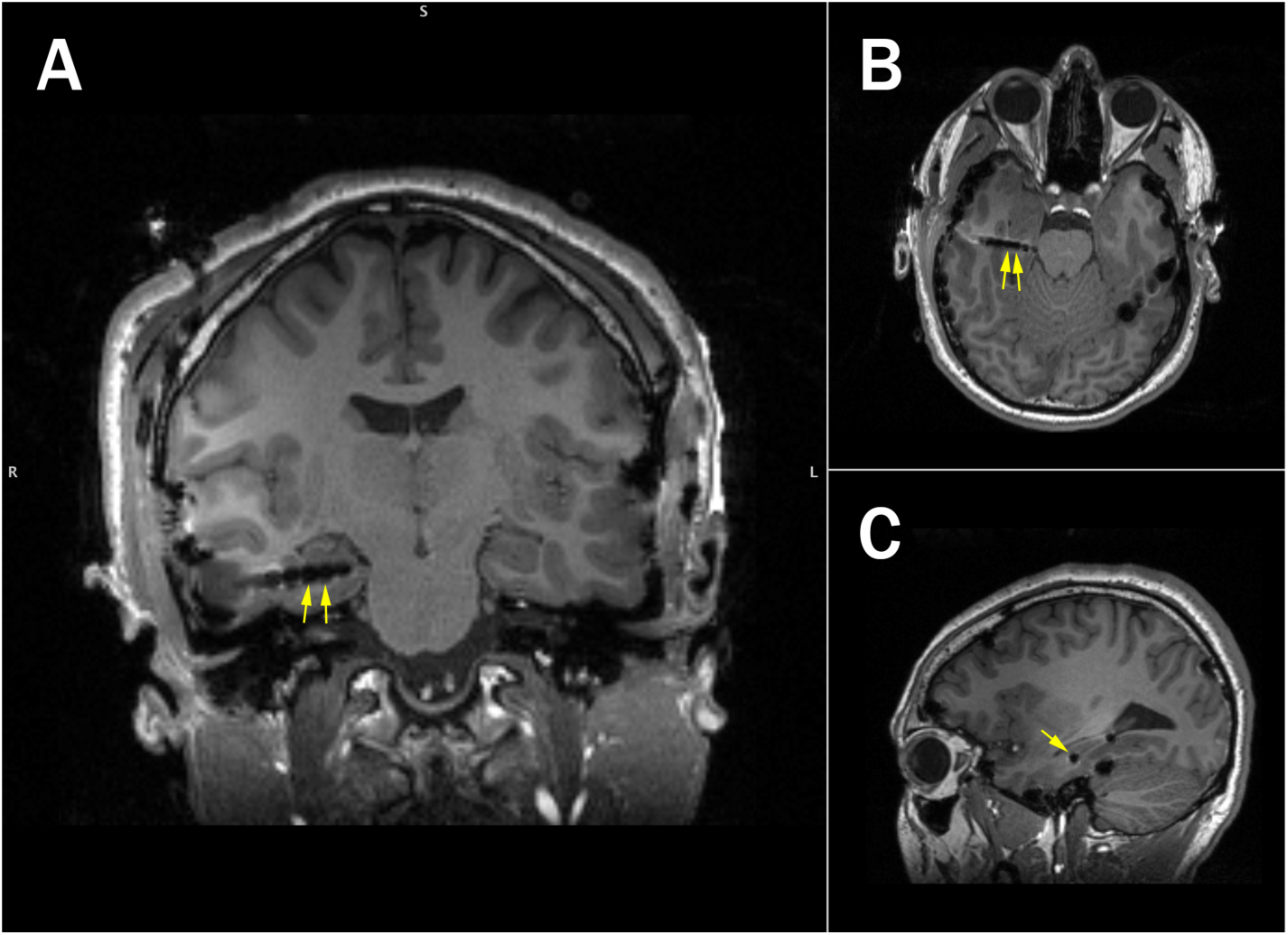
Electrode localization for an example surgical patient (S2). Location of selected electrodes is indicated with the yellow arrows (Right Anterior Temporal 03/04, A: Coronal, B: Axial, C: Sagittal).

### Task Design

#### Face-profession Association Task

A computer-based face-cued word recall task was used to test associative memory. A set of 240 high resolution color images of distinct human faces with neutral expression were taken from the Chicago Face Database (**Figure 5**). These images were divided in half to form two equivalent versions of the task, allowing for multiple testing sessions with the same patient. Each set of 120 faces were made up of an equal proportion of male and female White, Black, Hispanic, and Asian faces. Each face set was randomly paired with 120 emotionally neutral, single-word professions between 4-10 letters in length, selected from the US Bureau for Labor Statistics database.

**Figure 5.**
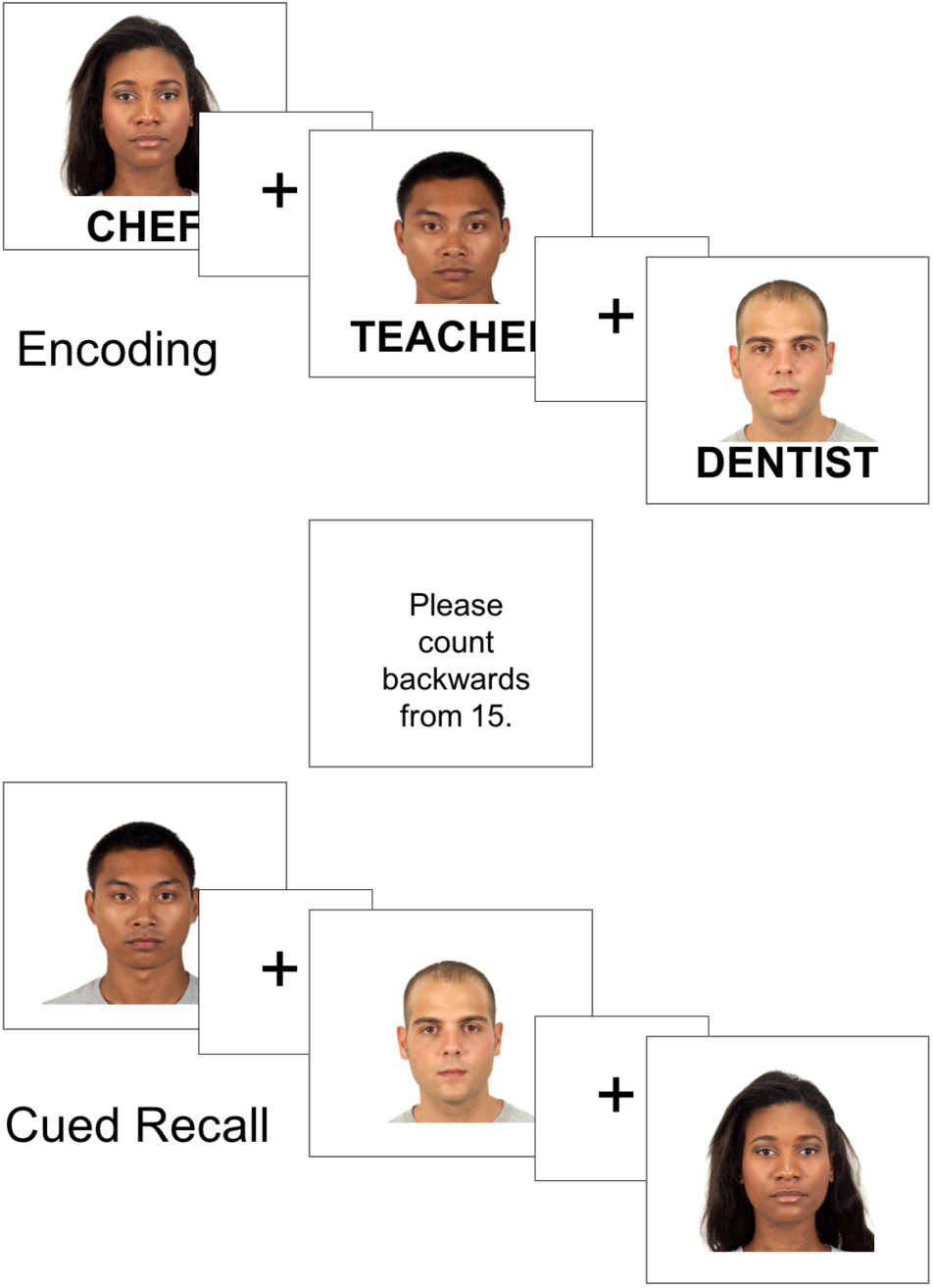
Face profession task. Patients were shown pairs of common professions and novel faces and asked to form an association. Following a distraction task, patients were cued with the previously seen faces and asked to freely recall the associated professions.

Given that performance varies between the inpatient and outpatient setting, the task was adjusted depending on patient performance in an initial practice set. To ensure attention and sensory processing of test stimuli, patients were instructed to read the profession aloud and make a mental association. Encoding blocks consisted of between 2-20 pairs, with each pair presented for 5000 ms with a 1000 ms inter-stimulus interval.

Individual encoding blocks were followed by a brief distraction task to prevent rehearsal and then a cued recall session during which patients were shown the faces from the encoding session, in a pseudo-randomized order, and asked to recall the associated profession. Responses were recorded and scored for offline analysis. This cued recall paradigm was chosen because it offers more experimental control than traditional free recall.

### Surgical and RNS System iEEG Data Analysis and Statistics

Hippocampal electrodes were selected according to electrode localization procedures described above, with expert validation by two independent reviewers (AL, DF). As iEEG signals recorded by the RNS System lead electrodes are bipolar, surgical iEEG recordings were re-referenced in a bipolar montage to facilitate comparison, using the electrode selection criteria described above. All candidate electrodes were inspected for signal quality by plotting the raw voltage tracings by trial. Electrodes were discarded based on high 60 Hz noise (likely due to poor contact impedance) and/or excessive interictal activity. Standard artifact rejection techniques, such as notch filtering for line noise and harmonics (60, 120, 180 Hz), detrending, and baseline correction were applied. In addition, EEG artifacts were removed by omitting trials where the raw signal exceeded 5 SD above the mean (across all trials per subject), as well as trials with excessive interictal epileptiform activity. For RNS System data, an additional data blanking routine was used to zero-out telemetry marker artifacts (“triggers”) delivered at stimulus onset (see **Figure 2**, Supplementary Figure 2). Finally, visual inspection of individual trials was used to exclude trials with excessive non-physiological noise. A summary of the trial rejection for each subject is in **Table 1**.

Analysis focused on identifying differences in the spectral-temporal features between successful and failed encoding trials. All trials from the selected electrodes were pooled for subsequent analyses. We performed time-frequency analyses on discrete task windows, including 1 s before and 3 s during encoding. EEG recordings were separated into trials of subsequently recalled and forgotten face-profession pairs. For each trial, gamma power was extracted using 8 semi-logarithmically spaced constant-Q Morlet wavelet filters (center frequencies between 65-100 Hz, 6 cycles) and z-scored relative a pre-stimulus baseline window (-0.5 to -0.05 s). To assess for differences between conditions (successful vs. failed encoding), a non-parametric permutation test was used to determine significance of any power changes between conditions and corrected for multiple comparisons using a cluster-based correction^27^.

In order to assess if successful versus failed encoding trials could be differentiated on a single trial basis, a Naïve Bayes classifier was used to decode the neural activity evoked between successful and failed encoding trials using 10-fold cross validation. Gamma power from the same 8 semi-logarithmically spaced filters calculated above, were used as the feature vector, and decoding accuracy was calculated at each time point in the 1-2s time period post stimulus onset where significant gamma power changes were observed in both RNS System and surgical patients. Training and classification was repeated 10 times and classification accuracy was averaged across repetitions (**Supplemental Figure S3**).

## AUTHOR CONTRIBUTIONS

AL, LD conceptualized the experiment. AL, SH, AS, CS designed the experiment. SH, AS, MF collected the data. AL, DF, PD supervised clinical aspects of the study. NH, TT designed the research accessories. NH, MF, TT provided technical support for RNS System. WD performed intracranial surgery and implanted the RNS System device. SH, AS, DF, AL analyzed the data. SH, AS, AL wrote the manuscript. SH, AS, NH, DF, PD, MF, LM, AF, MF, TT, OD, LD, AL provided critical revisions of the manuscript.

